# BIFO: A Biological Information Flow Ontology for Directed Propagation in Heterogeneous Biomedical Knowledge Graphs

**DOI:** 10.64898/2026.05.25.727605

**Authors:** Deanne M. Taylor, Taha Mohseni Ahooyi, Benjamin Stear, Yuanchao Zhang, Aditya Lahiri, J. Alan Simmons, Asif Chinwalla, Christopher Nemarich, Tiffany J. Callahan, Jonathan C. Silverstein

## Abstract

Biomedical knowledge graphs integrate heterogeneous data by connecting many entity types through many relationship types. Computational analyses that propagate signal across these graphs (random walks, diffusion, and message passing) implicitly assume that every traversable edge can carry a biological signal. In a heterogeneous KG this is rarely true: hierarchical, lexical, and purely statistical edges do not, by themselves, define an admissible directed state transformation, and traversing them propagates signal along paths that are not biologically meaningful. We present the Biological Information Flow Ontology (BIFO), a graph-agnostic specification of which directed transformations are biologically admissible for computable information flow. BIFO defines fourteen entity classes, a taxonomy of flow classes organized around the backbone `G+CH` →`RNA` →`P` →`PW` →`C` →`PH` →`DS`, a set of admissibility constraints, and a two-level CURIE mapping that can be applied without schema-specific code to any graph whose identifiers and predicates are resolvable through, or extendable to, the BIFO mapping tables. A four-step conditioning protocol converts a raw property graph into a conditioned propagation graph in which only admissible, direction-aware edges remain. We provide a reference implementation on the Data Distillery Knowledge Graph (DDKG); conditioning a cohort-independent, gene-anchored subgraph as a BIFO substrate of 33.6 million edges retained 23.7 million (70.7%) as BIFO-classified relationships, cleanly separating 13.3 million propagating mechanistic edges from 10.5 million retained-but-non-propagating observational associations, and confirming that pathway concepts are configured as scoring accumulation endpoints for BIFO-PPR pathway scoring. BIFO is an admissibility specification for computable propagation of signal over knowledge graphs. It is released as an open specification with versioned mapping tables and tooling, providing a reusable substrate for biologically interpretable, direction-aware analysis of biomedical knowledge graphs.

## Background & Summary

Biological systems are, at their core, organized by the flow of information: genetic sequence is transcribed and translated, signals propagate through molecular networks, and these transfers sustain the organization that distinguishes living systems from systems at equilibrium (Nicolis and Prigogine 1977). If biological function is fundamentally about how information moves through a system, a framework for analyzing biology should make that movement explicit and computable rather than leaving it implicit in static annotation. Knowledge graphs are a natural substrate for this view, because they encode complex biological systems as traversable topologies. The structure of a knowledge graph is a potentially computable map of how entities are connected, and therefore of how information could in principle move among them. To use such graphs as more than static catalogs of observations, that is, to compute over them and recover biological mechanisms through traversal, requires a principled account of which connections actually carry biological information and in which direction. The resulting problem is a computable semantics problem: how to determine, for each graph edge, whether traversal is biologically admissible.

Knowledge graphs (KGs) have become central tools for integrating heterogeneous biomedical data, linking genes, variants, regulatory elements, transcripts, proteins, pathways, cells, tissues, phenotypes, and diseases in a single structured representation. A growing class of analyses extracts biological meaning from these graphs by *propagating* signal across edges: random-walk and diffusion methods, personalized PageRank, and graph neural message passing. All of these methods share an implicit assumption: that an edge which can be traversed in a biomedical KG is an edge along which biological signal can meaningfully flow.

That assumption is usually false. A heterogeneous KG could mix mechanistic relationships, such as a transcription factor regulating a gene or an enzyme transforming a metabolite, with relationships that do not, by themselves, define an admissible directed state transformation: ontology hierarchy edges (is_a, part_of), identifier and lexical links, non-directional interaction evidence such as protein-protein contacts, and statistical associations such as co-expression correlations. Such edges may be biologically informative as annotation, context, or weighting, but they do not specify a direction of biological information transfer. Propagating across them can diffuse signal along paths that are not admissible under the specified propagation semantics and can inflate connectivity significance between entities related only by annotation convenience. Equally problematic, most meaningful relationships in biological systems are directional: information commonly flows from a regulatory element to a transcript, but the reverse requires special regulatory cases in directional feedback loops. Methods that traverse edges symmetrically discard these cases of directionality and the constraints they impose.

Existing frameworks do not close this gap. Reference ontologies such as the Gene Ontology and the Human Phenotype Ontology provide rich semantic representations but do not specify which relationships may propagate biological signal (Ashburner et al. 2000; Köhler et al. 2021). Mechanistic modeling standards such as SBML encode quantitative models but are not designed as graph-agnostic conditioning layers over heterogeneous KGs (Hucka et al. 2015). Causal-assertion frameworks such as GO-CAM and the Biological Expression Language describe curated causal relationships but do not generalize to arbitrary heterogeneous graphs (Slater 2014; Thomas et al. 2019). Graph machine-learning approaches learn propagation behavior from data rather than enforcing biologically interpretable constraints, sacrificing transparency for flexibility. Schema standards such as the Biolink Model harmonize entity and predicate *types* across graphs but do not assert which typed edges are admissible for biological information *propagation* (Unni et al. 2022). The missing layer is therefore not another biological ontology of entities, but a computable admissibility layer over graph assertions.

BIFO addresses this missing layer directly. By analogy to communication systems, where transmission depends on admissible channels, BIFO treats graph edges as candidate channels for biological state propagation, defining admissible *biological information flow* independently of the underlying graph schema, identifier system, or database (Shannon 1948). The analogy is operational rather than physical: BIFO specifies which graph assertions may be traversed by propagation algorithms, not the thermodynamic information content of living systems. The asymmetry introduced by admissible directional flows constrains traversal to semantically valid biological paths, adding information by restricting the reachable state space. BIFO comprises four components: a set of entity classes representing the organizational layers of biology; a taxonomy of flow classes defining admissible directed transformations; a set of admissibility constraints (Fig 1) and a two-level CURIE mapping that makes the ontology applicable to any standards-compliant graph. A four-step conditioning protocol applies these components to convert a raw property graph into a conditioned propagation graph. We describe each component and demonstrate the framework through a reference implementation on the Data Distillery Knowledge Graph.

**Figure 1.**
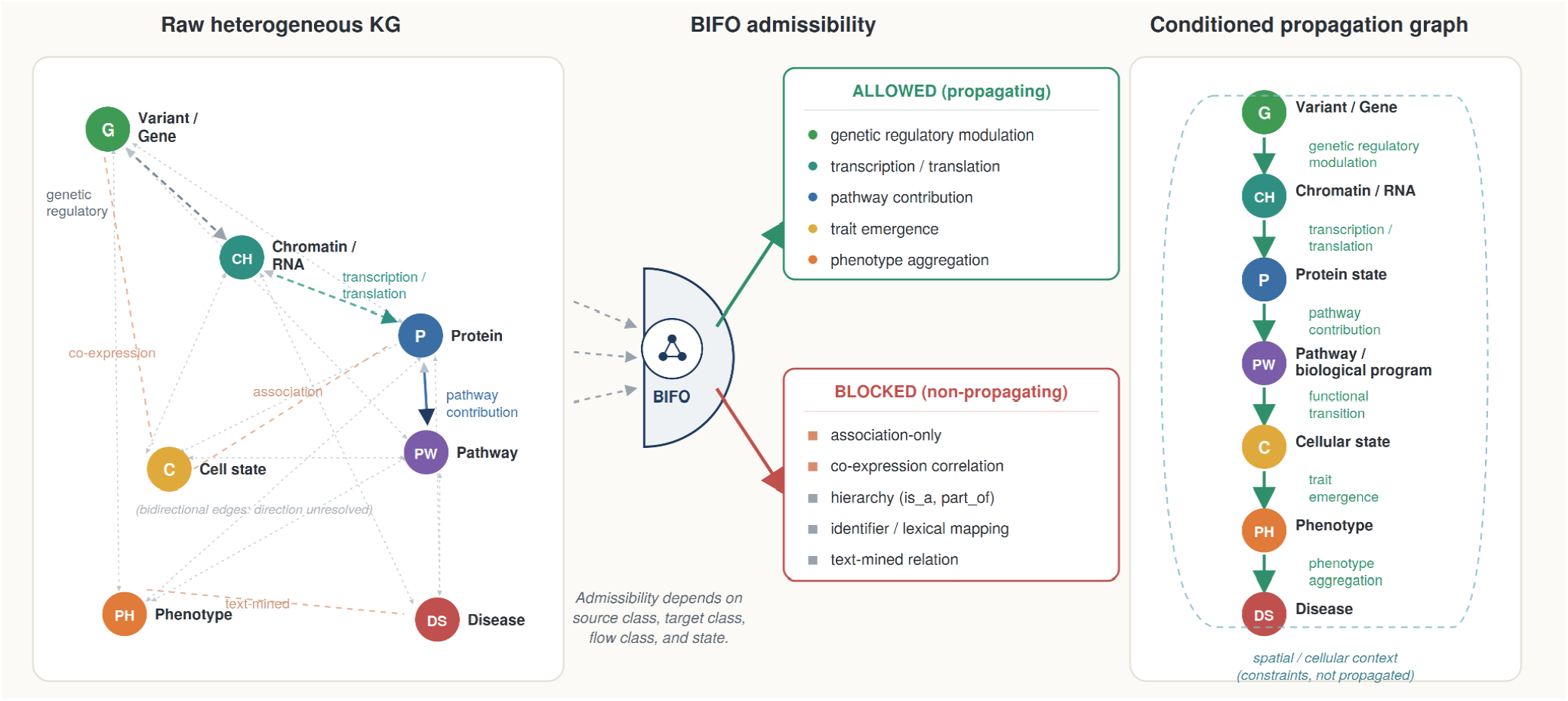
The BIFO admissibility operator conditions a raw heterogeneous knowledge graph into a directed propagation substrate. **Left:** a raw heterogeneous biomedical knowledge graph connects biological entities (genes and variants, chromatin and RNA, proteins, pathways, cells, phenotypes, and diseases) through many relationship types. Most edges are stored bidirectionally and mix genuine mechanistic relationships with non-mechanistic ones (ontology hierarchy, identifier and lexical links, statistical associations such as co-expression, and curated edges that connect entities not adjacent in information-flow terms, for example direct gene-to-pathway links). In this raw state the admissible direction of flow is unresolved. **Center**: the BIFO admissibility operator evaluates each candidate edge using its source entity class, target entity class, flow class, and state, and partitions edges into those that are admissible for biological information flow (propagating) and those that are retained but non-propagating (observational associations, hierarchy, and text-mined relations). Identifier or lexical links are usually dropped. **Right:** the resulting conditioned propagation graph retains only admissible, direction-aware edges along the information-flow backbone (G/variant to chromatin and RNA to protein to pathway to cellular state to phenotype to disease). Spatial and cellular context (dashed envelope) constrains admissibility but is not itself propagated.

BIFO is complementary to other structural ontologies. Anatomy, cell, and process ontologies such as UBERON, the Cell Ontology, and the Gene Ontology define what biological entities are and how they are organized, but they do not, by design, assert how information flows along their structures: that is not their purpose (Mungall et al. 2012; Osumi-Sutherland 2017). BIFO supplies precisely this missing layer, the directionality and admissibility of information flow over structures these resources already define, and users of a knowledge graph can use such ontologies for biological structure and BIFO for flow semantics at the same time.

## Methods

### Entity classes

BIFO is an operational ontology specification for propagation over biomedical KGs: its classes and relations define admissible computational roles. BIFO defines fourteen entity classes representing the major organizational layers of biology, each carrying an internal state appropriate to its role (**Table 2**). A BIFO class is the role an entity plays in a specified propagation context; a source graph may map the same underlying biological entity differently under different predicates or contexts. The classes organize into operational roles, into an informational state space: `G` (genetic sequence and variation), `CH` (chromatin state), `RNA` (transcripts), `P` (proteins), `CM` (macromolecular complexes), `PW` (pathways and programs), `C` (cellular configuration), `PH` (phenotype), and `DS` (disease); and a resource and physical state space: `SM` (small molecules), `ION` (ionic and electrochemical state), and `MECH` (mechanical state), together with `S` (spatial context) and `X` (external perturbations). Informational flow drives system behavior, while resource and physical states constrain admissible transitions. The classes are organized around a primary information-flow backbone, `G+CH` →`RNA` →`P(state)` →`PW` →`C` →`PH` →`DS`, which traces the path from genetic sequence and chromatin state through transcription and translation to pathway activity, cellular configuration, phenotype, and disease (**Figure 1**). When state metadata are absent, BIFO applies only the flow rules that can be evaluated from endpoint class and predicate; state-dependent refinements are applied only when the source graph provides the required state information.

**Table 1.** Positioning of BIFO relative to existing ontologies, standards, and graph-analysis approaches. Among the frameworks compared here, BIFO is the only one whose primary purpose is to specify admissible typed, directed edges for biological information propagation over heterogeneous KGs, and others are complementary layers that BIFO builds on or runs alongside.

| Resource /approach | What it provides | Asserts which typed edges are admissible for directed biological propagation? | Relation to BIFO |
| --- | --- | --- | --- |
| GO, HPO (reference ontologies) | Rich semantic representation of processes, functions, phenotypes | No | Complementary; BIFO adds the flow layer over the structures they define |
| Relation Ontology (RO) | Standard relation types with formal semantics | RO provides formal relation semantics that BIFO can reuse, but RO does not by itself define which graph assertions should be traversed by propagation algorithms. | Used by BIFO at Level 2 to resolve predicates to flow classes |
| Biolink Model | Harmonized entity and predicate types across graphs | No (harmonizes categories and predicates but does not define a propagation-admissibility operator. | Compatible; BIFO adds admissibility on top of harmonized types |
| GO-CAM, BEL (causal assertion) | Curated causal relationships | Partially, for curated assertions; does not generalize to arbitrary heterogeneous graphs | Narrower scope; BIFO is graph-agnostic and operates on any standards-compliant KG |
| SBML, SBGN (mechanistic models) | Quantitative kinetic / biochemical network models | No (models dynamics, not graph-traversal admissibility) | Different layer; BIFO conditions graphs rather than simulating kinetics |
| Graph ML / GNNs | Propagation behavior learned from data | No (learned, not specified; not biologically interpretable by construction) | BIFO supplies an interpretable, specified admissibility layer such methods can run on |
| BIFO (this work) | Graph-agnostic specification of admissible directed information flow | Yes | Conditioning layer |

**Table 2.**
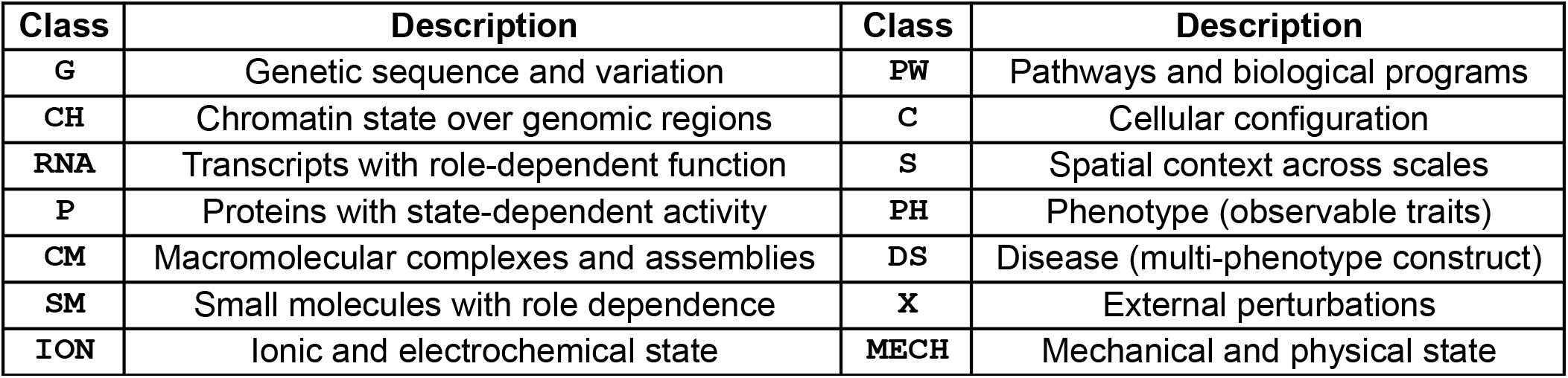
BIFO entity classes. Each class carries an internal state (e.g., protein PTM state and activity; chromatin accessibility and methylation) used to evaluate flow admissibility.

| Class | Description | Class | Description |
| --- | --- | --- | --- |
| G | Genetic sequence and variation | PW | Pathways and biological programs |
| CH | Chromatin state over genomic regions | C | Cellular configuration |
| RNA | Transcripts with role-dependent function | S | Spatial context across scales |
| P | Proteins with state-dependent activity | PH | Phenotype (observable traits) |
| CM | Macromolecular complexes and assemblies | DS | Disease (multi-phenotype construct) |
| SM | Small molecules with role dependence | X | External perturbations |
| ION | Ionic and electrochemical state | MECH | Mechanical and physical state |

**Table 3.** BIFO conditioning of the cohort-independent all-HGNC DDKG subgraph (10.6M nodes, 33.6M edges). A majority of concept edges are retained as admissible; dropped edges are dominated by unresolved-endpoint provenance/terminology nodes. Observational associations are retained but flagged non-propagating.

| Conditioning metric | Value |
| --- | --- |
| Nodes total | 10,646,735 |
| – Resolved to BIFO entity class | 6,152,488 (57.8%) |
| Concept-concept edges | 33,581,300 |
| Kept after conditioning | 23,739,700 (70.7%) |
| – Propagating | 13,264,410 |
| – Non-propagating (observational, retained) | 10,475,290 |
| Dropped: unresolved entity | 9,355,096 |
| Dropped: non-flow classification | 321,814 |
| Dropped: unmapped predicate | 164,690 |
| Pathway nodes: inbound propagating edges | 1,208,983 |
| Pathway nodes: outbound propagating edges | 0 |

**Table 4.** Distribution of the 13,264,410 propagating edges across BIFO flow classes in the conditioned all-HGNC DDKG subgraph as substrate. Signal Transduction (which absorbs the regulatory predicate families via entity-pair-dependent resolution) and Transcription dominate; Pathway Contribution edges bridge molecular entities to pathway programs. Perturbational Effect comprises the admissible small-molecule-to-gene targeting edges; statistical chemical/drug correlation predicates and non-admissible reverse orientations are retained as observational and excluded from this propagating distribution.

| Propagating flow class | Edges |
| --- | --- |
| Signal Transduction | 7,870,577 |
| Transcription | 3,582,746 |
| Perturbational Effect | 669,486 |
| Pathway Contribution | 552,656 |
| Signal Termination | 512,433 |
| Translation | 48,210 |
| Genetic Regulatory Modulation | 26,908 |
| Directed Protein Interaction | 1,245 |
| Biochemical Transformation | 137 |
| Other (Molecular→Phenotype, Complex Formation) | 12 |

**Table 5.**
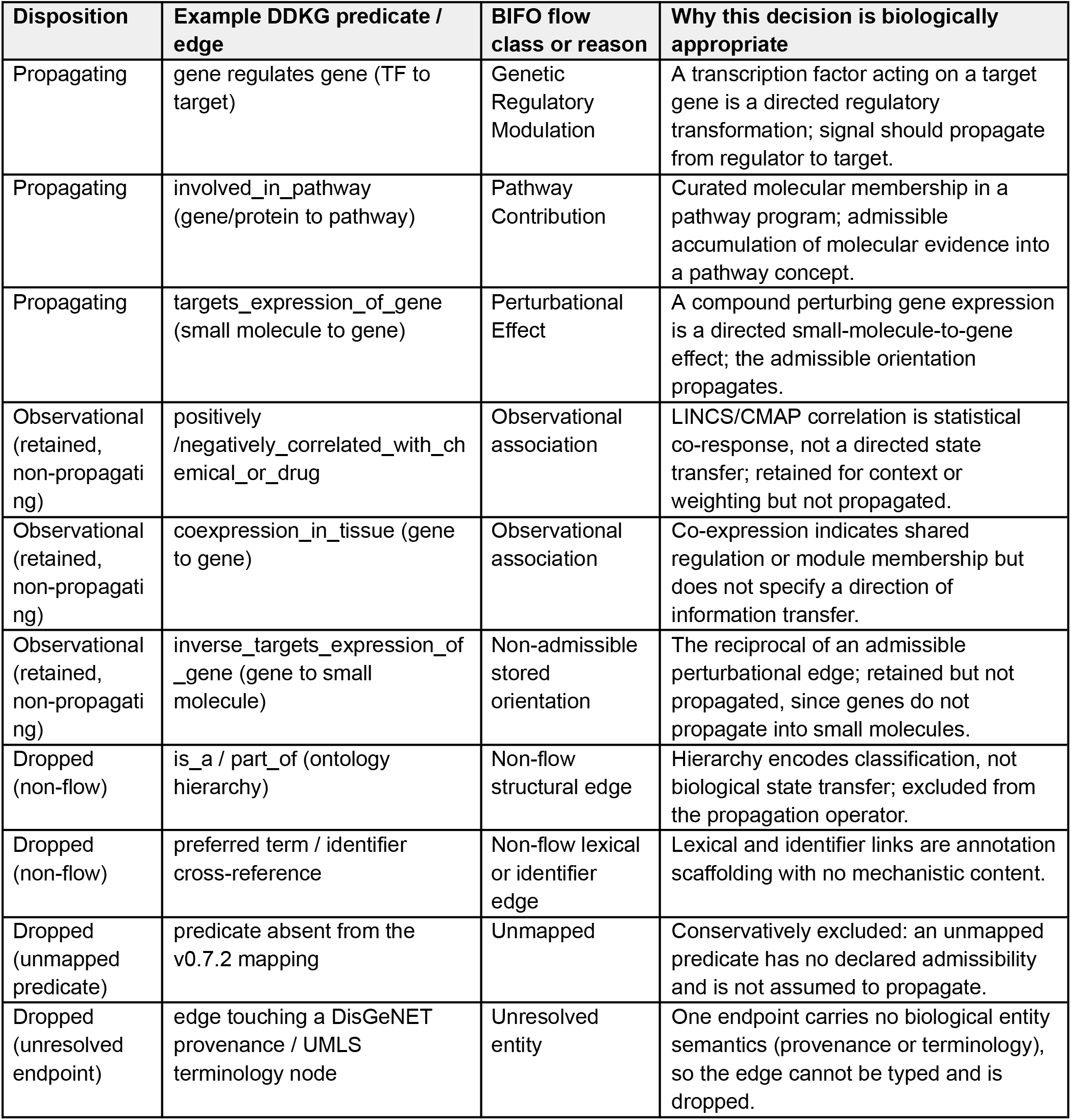
Representative edges in each conditioning disposition, with the biological rationale for the BIFO decision. Examples are drawn from the DDKG reference implementation. This audit-style table illustrates that conditioning decisions are biologically motivated at the level of individual predicates, not only large in aggregate

| Disposition | Example DDKG predicate / edge | BIFO flow class or reason | Why this decision is biologically appropriate |
| --- | --- | --- | --- |
| Propagating | gene regulates gene (TF to target) | Genetic Regulatory Modulation | A transcription factor acting on a target gene is a directed regulatory transformation; signal should propagate from regulator to target. |
| Propagating | involved_in_pathway (gene/protein to pathway) | Pathway Contribution | Curated molecular membership in a pathway program; admissible accumulation of molecular evidence into a pathway concept. |
| Propagating | targets_expression_of_gene (small molecule to gene) | Perturbational Effect | A compound perturbing gene expression is a directed small-molecule-to-gene effect; the admissible orientation propagates. |
| Observational (retained, non-propagating) | positively /negatively_correlated_with_chemical_or_drug | Observational association | LINCS/CMAP correlation is statistical co-response, not a directed state transfer; retained for context or weighting but not propagated. |
| Observational (retained, non-propagating) | coexpression_in_tissue (gene to gene) | Observational association | Co-expression indicates shared regulation or module membership but does not specify a direction of information transfer. |
| Observational (retained, non-propagating) | inverse_targets_expression_of_gene (gene to small molecule) | Non-admissible stored orientation | The reciprocal of an admissible perturbational edge; retained but not propagated, since genes do not propagate into small molecules. |
| Dropped (non-flow) | is_a / part_of (ontology hierarchy) | Non-flow structural edge | Hierarchy encodes classification, not biological state transfer; excluded from the propagation operator. |
| Dropped (non-flow) | preferred term / identifier cross-reference | Non-flow lexical or identifier edge | Lexical and identifier links are annotation scaffolding with no mechanistic content. |
| Dropped (unmapped predicate) | predicate absent from the v0.7.2 mapping | Unmapped | Conservatively excluded: an unmapped predicate has no declared admissibility and is not assumed to propagate. |
| Dropped (unresolved endpoint) | edge touching a DisGeNET provenance / UMLS terminology node | Unresolved entity | One endpoint carries no biological entity semantics (provenance or terminology), so the edge cannot be typed and is dropped. |

### Flow classes

Flow classes define the admissible directed transformations between entities. They are organized into biologically coherent groups: central-dogma transformations (Transcription, `G+CH` →`RNA` ; Translation, `RNA` →`P`); chromatin regulation; RNA-mediated regulation; protein and complex signaling (including Signal Transduction, `P(state)` →`P(state′),` and Pathway Contribution, `P or CM` →`PW`); genetic modulation of transcription; small-molecule, redox, and electrochemical transformations; spatial and transport processes; cell-cell communication; cellular state and progression; mechanotransduction; and phenotype/disease emergence (`C` →`PH),(PH` →`DS)`. The flow `PH` →`DS` represents evidence accumulation from observed phenotypes toward disease concepts for scoring or interpretation; it is not intended to assert that phenotypes mechanistically cause diseases. In binary graph implementations, the flow `G+CH` →`RNA` can also be represented as `G` →`RNA` conditioned by chromatin-state context, or as a reified regulatory assertion when the source graph represents chromatin state explicitly.

Each flow is defined by its source and target entity classes and a direction reflecting biological causality. Critically, a separate set of observational relationships (single-cell clustering assignments, accessibility correlations, co-occurrence, and topological proximity) are explicitly designated as non-propagating: they may inform weighting or constraints but must not carry biological state during propagation. This mechanistic/observational definition, in practice, is enforced during graph conditioning. Non-directional interaction evidence is retained as observational unless the predicate and endpoint state information support a directed flow class such as signal transduction, complex formation, signal termination, or biochemical transformation; co-expression may support weighting along a flow, context for a flow, or module definition, but does not by itself usually specify a direction of biological information transfer. Local extensions may promote otherwise non-directional interaction evidence to a propagating class only when the extension documents the biological direction, endpoint roles, and state/context assumptions, for example by defining transcription factor binding processes on otherwise non-directional protein-protein interaction data.

### Admissibility constraints and the conditioning operator

All flows are governed by admissibility constraints: directionality (forward and reverse of a transformation are distinct flows with independent conditions), state dependence, spatial compatibility, temporal progression, and the exclusion of non-flow edges (hierarchy, identifier/lexical, and purely statistical relationships). Formally, for an input property graph **G**=(V,E) in which each edge carries a predicate and endpoint entity classes, BIFO conditioning defines a constraint operator **C** that yields a conditioned propagation graph **G**_**C**_=(V,E_C_). retaining an edge if and only if its predicate maps to an admissible flow class and both endpoints resolve to BIFO entity classes. BIFO conditioning then partitions edges into three dispositions: propagating, retained non-propagating observational, and dropped. Therefore, the propagation substrate is the propagating subset; the conditioned audit graph may also retain observational edges flagged as non-propagating. The operator is graph-agnostic: it depends only on the predicate-to-flow mapping and entity resolution, not on graph topology.

### KG-agnostic CURIE mapping

BIFO achieves graph independence through a two-level CURIE mapping. Level 1 maps node identifiers (CURIEs) from standard biomedical ontologies to entity classes, for example HGNC (Seal et al. 2023) and Ensembl (Harrison et al. 2024) gene identifiers to `G` ; UniProtKB(UniProt Consortium 2025) to `P` ; Reactome (Joshi-Tope et al. 2005), GO biological process, and WikiPathways (Pico et al. 2008) to `PW` ; MONDO (Shefchek et al. 2020) and OMIM (Amberger et al. 2018) to `DS`. Level 2 maps relationship predicates, including Relation Ontology terms^8^, to flow classes, with entity-type-dependent rules resolving predicates that can express multiple flows (Smith et al. 2005). Both mappings are normative and KG-agnostic, so any graph whose nodes and edges carry resolvable CURIEs can be conditioned without schema-specific code. A graph’s BIFO compatibility can be quantified as the fraction of its nodes and edges that resolve through these tables.

### Conditioning protocol and provenance

BIFO, as a set of rules to transform a biomedical knowledge graph, can be applied by a logical protocol. The protocol (**Figure 2**) assigns each node an entity class (Step 1); assigns each edge a flow class and removes non-admissible edges (Step 2); annotates nodes and edges with state and spatial context (Step 3); and annotates retained edges with confidence and provenance (Step 4). The default evidence tiers are coarse ordinal categories, not calibrated probabilities, and implementations may override them with source- or instance-level evidence metadata when available. For multi-hop composite flows, path confidence is the lowest ordinal evidence/confidence tier along the path (weakest-link principle), typed at the lowest evidence tier present. Where a predicate type has mixed evidence quality across instances, the protocol applies the weaker classification, a conservative choice that prevents high-confidence mechanistic relationships from being diluted by observational instances of the same predicate.

**Figure 2.**
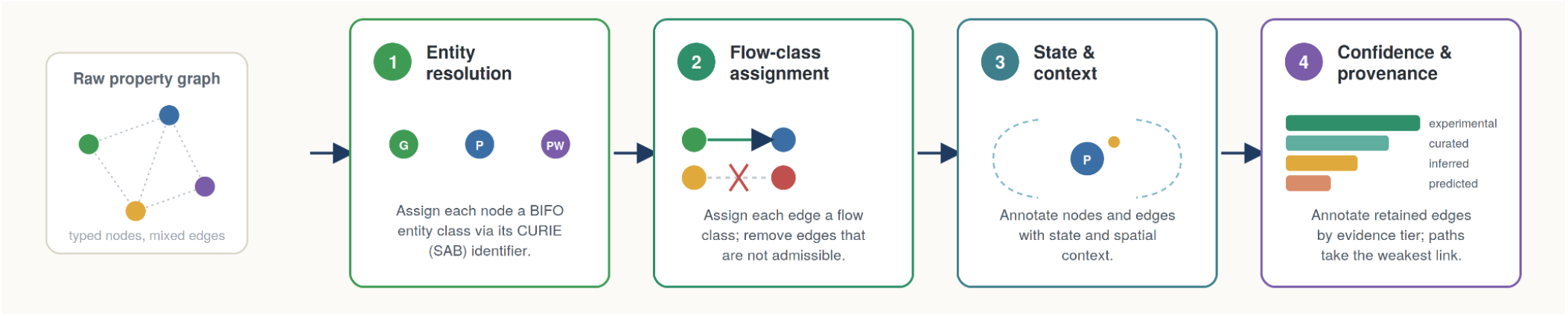
The four-step BIFO conditioning protocol. A raw heterogeneous property graph is converted into a conditioned propagation graph in four steps. Step 1 (entity resolution): each node is assigned a BIFO entity class from its CURIE or source-authority (SAB) identifier. Step 2 (flow-class assignment): each edge is assigned a flow class using its endpoint entity classes and predicate, and edges that do not map to an admissible flow are removed. Step 3 (state and context): retained nodes and edges are annotated with internal state and spatial or cellular context, which constrain admissibility without themselves propagating. Step 4 (confidence and provenance): retained edges are annotated by evidence tier (experimental, curated, inferred, predicted); for multi-hop composite flows, path confidence follows the weakest link and is typed at the lowest evidence tier present. The output is a conditioned propagation graph of admissible, direction-aware edges, with non-propagating observational associations retained and flagged.

### Evidence use in BIFO

BIFO by design separates propagation semantics from evidence semantics. Flow admissibility is determined by endpoint entity classes, predicate-to-flow mapping, directionality, state, and context; it is not inferred from strength of evidence. Evidence metadata may be used by BIFO-compatible implementations to qualify retained relationships, support auditability, and enable downstream filtering or weighting. Propagating edges may therefore carry both a flow class and confidence/provenance metadata, while observational relationships may be retained as evidence, context, constraints, or candidate weights, but do not themselves carry biological state during propagation. When an implementation reports confidence for composite or multi-hop paths, the BIFO specification recommends a conservative weakest-link convention, so a composite path is never treated as more reliable than its least-supported edge.

## Data Records

### Specification

BIFO is distributed as a normative specification (entity classes and state definitions; the two-level CURIE mapping; flow-class definitions; admissibility constraints; a four-step conditioning protocol; and confidence/provenance annotation) together with a reference DDKG implementation guide and a versioned predicate-to-flow mapping YAML file. The reference conditioning run is reproducible from the deposited data records: gene-anchored extraction queries defining the all-HGNC subgraph (substrate); a versioned mapping used to condition it, and the resulting coverage report; per-flow-class breakdown; and conditioned (kept) and dropped edge lists. Tissue-specific co-expression predicates from GTEx, which are classified observational, are excluded at query extraction to limit the calculational size for BIFO; all other observational edges are retained and flagged non-propagating during conditioning. The specification is released under CC BY 4.0 and the accompanying mapping and tooling under the MIT License. The current release is BIFO v0.2.2 with DDKG mapping v0.7.2. The specification, mapping, and data records are available at https://github.com/TaylorResearchLab/BIFO and archived at Zenodo (https://doi.org/10.5281/zenodo.20370680). The deposited data records comprise the gene-anchored extraction queries, the versioned mapping, the conditioning code, and the coverage and per-flow-class reports; the full conditioned (kept) and dropped edge lists are regenerable from these inputs as described in the repository.

### FAIR alignment

BIFO is designed as a reusable, graph-agnostic specification rather than a KG-specific implementation. Findability and accessibility are supported by versioned releases, GitHub distribution, Zenodo archiving, and explicit licensing. Interoperability is supported by CURIE-based entity resolution, predicate-to-flow mappings, and use of community biomedical identifiers. Reusability is supported by the deposited mapping tables, conditioning code, coverage reports, flow-class summaries, and regenerable kept/dropped edge records. These features make BIFO FAIR-aligned while preserving the distinction between the normative flow specification and graph-specific implementation choices. The BIFO specification, mappings, and tooling are openly available; regeneration of the DDKG reference conditioning run requires access to the underlying DDKG release, including UMLS licensing requirements (see Technical Validation).

### Technical Validation

We validated BIFO through a reference implementation on a defined substrate: a subgraph from the Data Distillery Knowledge Graph (DDKG), built on the Unified Biomedical Knowledge Graph (UBKG) and Petagraph infrastructure, instantiated on Neo4j 5.26.25 community edition. The reference build was the DDKG version 2025-JUL-28 (available at https://ubkg-downloads.xconsortia.org/; download requires a free UMLS license key). DDKG is an appropriate reference graph because it integrates summarized Common Fund data within a UBKG-based graph and links multiple expertly curated data sources across Common Fund data coordinating centers, supported by more than 180 ontologies and standards(Stear et al. 2024; Ahooyi et al. 2025). The subgraph was extracted as the undirected one-hop Concept-layer neighborhood of all 43,838 HGNC gene concepts in DDKG, combined with the complete gene-to-pathway membership universe for the enabled pathway source, MSigDB v 7.4 (selecting Hallmark and C2.CP minus CGP and KEGG)(Liberzon et al. 2011). This subgraph is defined by gene identity and graph version, and no study cohort, case-control label, or study phenotype enters its construction. Therefore, the resulting coverage statistics are properties of the BIFO specification applied to this DDKG build, not of any particular biological cohort. The subgraph comprised 10,646,735 nodes and 33,592,072 edges, of which 33,581,300 were concept-concept relationships subject to conditioning. DDKG concept nodes were resolved to BIFO entity classes via their source-authority identifiers using a versioned mapping comprising 252 predicate-to-flow entries, 96 explicit non-flow exclusions, and 46 observational definitions across five classification tiers. Overall, 6,152,488 of 10,646,735 nodes, or 57.8%, resolved to a BIFO entity class. The unresolved remainder consisted predominantly of provenance and terminology vocabularies, including DisGeNET source records and UMLS Metathesaurus concepts, that do not carry biological entity semantics for BIFO propagation and are excluded by design.

Applying the conditioning operator retained 23,739,700 of 33,581,300 concept-concept edges, or 70.7%, as BIFO-classified relationships. Of these, 13,264,410 were propagating edges and 10,475,290 were retained as non-propagating observational associations, including statistical correlation predicates and edges whose stored direction does not satisfy an independently admissible BIFO flow rule. Dropped edges fell into interpretable categories: 9,355,096 had an unresolved endpoint entity, overwhelmingly from the provenance and terminology vocabularies noted above; 321,814 were explicit non-flow structural edges; and 164,690 carried predicates not mapping to any flow class. Among propagating edges, the dominant flow classes were Signal Transduction, Transcription, Perturbational Effect, Pathway Contribution, and Signal Termination, with Translation, Genetic Regulatory Modulation, and other classes in the long tail. In this DDKG pathway-scoring projection, the reference mapping configures pathway concepts as terminal scoring nodes: they received 1,208,983 inbound propagating Pathway Contribution edges and emitted no outbound propagating edges. Thus, for this scoring projection, pathway concepts accumulate molecular evidence for pathway-level analysis rather than redistributing signal back to genes or other annotations.

This terminal scoring behavior is a property of the DDKG pathway-scoring projection, not a general BIFO axiom. In the BIFO ontology, `PW` denotes pathway or program state and may participate in downstream flows toward cellular configuration, phenotype, and disease; other BIFO-conditioned graphs may therefore enable `PW` →`C` , `PW` →`PH,` or `PW` →`DS` transitions. In the pathway-scoring projection, however, Pathway Contribution edges represent admissible accumulation of molecular evidence into pathway or program concepts for scoring. They do not assert that a gene dynamically causes a physically instantiated pathway state. The conditioned graph thereby separates a directional mechanistic propagation backbone from retained-but-non-propagating observational associations, demonstrating that BIFO conditioning can create mechanistic, observational, propagating, and accumulating distinctions on a large heterogeneous biomedical graph. This technical validation evaluates computability, coverage, auditability, and consistency of the BIFO specification on a large KG substrate; it does not claim independent biological validation of every predicate-to-flow assignment.

### Usage Notes

Any property graph satisfying three requirements is BIFO-compatible: 1. nodes represent biological entities carrying typed internal state; 2. edges carry sufficient semantic information to be assigned to BIFO flow classes; and 3. the graph supports provenance metadata sufficient to evaluate evidence type. BIFO does not require bidirectional edges, UMLS structure, or source-tagged provenance (Bodenreider 2004)^1^; a graph meeting the requirements only partially is BIFO-compatible for the subset of flow classes whose requirements are met. The conditioned propagation graph **G**_**C**_ is intended as the substrate for direction-aware propagation and traversal analyses; quantitative pathway-analysis methods built on BIFO-conditioned graphs are described separately (BIFO-PPR, in preparation). Because the conditioning operator depends only on the CURIE and predicate mappings, BIFO-compatible implementations may be applied to any standards-compliant biomedical KG, and KG-specific implementation guides would need to document any local extensions to the normative mapping tables.

Biological information flow can also be reified within a knowledge graph, with context represented as nodes that a flow passes through, for example tissue or cell-type nodes drawn from UBERON or the Cell Ontology, or reified expression and perturbation concepts that link an entity to the context in which a measurement was made. BIFO by design is agnostic to how an implementor chooses to propagate information over such a reified structure. The flow propagation strategy is a decision for the implementor and the schema of their graph, not something BIFO needs to prescribe. In particular, context-confinement, defined as restricting propagation so that signal flows only within a chosen biological context (such as within a single tissue, cell type, or subcellular compartment) is a useful condition an implementor may choose to impose on their own graph schema. It is important to be able to express, but it is neither required by BIFO nor uniform across graphs; whether and how it is applied depends entirely on the schema and on the way BIFO is being implemented by the user.

BIFO-compatible implementations may be used to define biological information flow across a range of contexts, provided those contexts are biologically sensible as translated to the system. The DDKG subgraph implementation we discuss here is, in effect, a human knowledge graph augmented with mouse model-organism evidence (orthology and phenotype mappings) interpreted in service of understanding human biology. These cross-organism relationships are not used as mechanisms but are used as evidence: biological information does not flow between organisms, and these mappings are accordingly treated as non-propagating associations rather than admissible flow. Applying BIFO to a different organism is therefore a matter of instantiating its flow rules over that organism’s own knowledge graph, not of joining organisms within a single flow space.

### Limitations

BIFO has several deliberate limitations. First, BIFO is an admissibility model, and should not be considered a kinetic or quantitative causal model: it specifies which directed transformations may carry propagated signal for purposes such as scoring hypotheses, but does not currently support modeling rates, magnitudes, or mechanisms of biological dynamics. Second, flow admissibility depends on the available predicate semantics and endpoint typing by the source knowledge graph; a predicate that is ambiguous or weakly specified in the source graph constrains what BIFO implementations can assert about it. Third, ambiguous predicates are conservatively classified at the weaker evidence tier, which favors specificity over sensitivity and may exclude relationships that are genuinely mechanistic in some instances. Fourth, context confinement (restricting information propagation to/within a tissue, cell type, or compartment) is implementer-defined and depends on the schema of the target graph; BIFO supports it but does not mandate it. Fifth, coverage depends on CURIE and predicate resolvability: nodes whose identifiers do not resolve to CURIEs from predominantly provenance and terminology vocabularies are intentionally excluded, so coverage statistics reflect resolvable biological content rather than the entire graph. Finally, BIFO does not determine which propagation algorithm is best; it defines the substrate on which propagation, diffusion, and message-passing methods can be biologically constrained, and the choice and evaluation of a specific method is left to downstream work.

## Code Availability

The BIFO specification, documentation (entity classes, CURIE mapping, flow classes, admissibility constraints, the conditioning protocol, and confidence and provenance annotation), the DDKG implementation guide, and the versioned predicate-to-flow mapping file (bifo_mapping_ddkg.yaml v0.7.2) are available at https://github.com/TaylorResearchLab/BIFO and archived at Zenodo (https://doi.org/10.5281/zenodo.20370680). Specification text is licensed CC BY 4.0; accompanying tooling and mapping files are licensed MIT. The current release is BIFO v0.2.2 with DDKG mapping v0.7.2.

